# Decoupling the correlation between cytotoxic and exhausted T lymphocyte transcriptomic signatures enhances melanoma immunotherapy response prediction from tumor expression

**DOI:** 10.1101/2023.01.17.524482

**Authors:** Binbin Wang, Kun Wang, Peng Jiang, Eytan Ruppin

## Abstract

**Background:** Cytotoxic T lymphocytes (CTL) play a crucial role in anti-cancer immunity. Progression of CTL to terminal exhausted T lymphocytes (ETL) that overexpress inhibitory receptors can substantially decrease effector cytokines production and diminish cytolytic activity and terminal exhausted T cell cannot be reprogrammed by ICIs in tumor microenvironment (TME). However, while the activity levels of CTL and ETL are considered important determinants of immune checkpoint inhibitors (ICIs) response, it has been repeatedly observed that their predictive power of the latter is quite limited. Studying this conundrum on a large scale across the TCGA cohort, we find that ETL and CTL activity (estimated based on conventional gene signatures in the bulk tumor expression) is strongly positively correlated in most cancer types. We hypothesized that the limited predictive power of CTL activity might result from the high concordance of CTL and ETL activities, which mutually cancels out their individual antagonistic effects on ICI response.

**Methods:** Consequently, we have set out to identify a set of genes whose expression identifies a subset of patients where the CTL and ETL correlation is diminished, such that the association between these CD8+ T cell states and ICIs response is enhanced.

**Results:** Analyzing TCGA melanoma bulk gene expression, we identified a set of genes whose over-expression markedly diminishes the CTL and ETL correlation, termed a *decoupling signature (DS)*. Reassuringly, we first find that the correlation between ETL and CTL activities is indeed markedly lower across high scoring DS patients than that observed across low scoring DS patients in numerous independent melanoma ICIs cohorts. Second, indeed, this successful decoupling increases the power of CTL activity in predicting ICIs response in high DS scoring patients. We show that the resulting prediction accuracy is superior to other state-of-art ICI predictive transcriptomic signatures.

**Conclusion:** The new decoupling score boosts the power of CTL activity in predicting ICIs response in melanoma from the tumor bulk expression. Its use enables a two-step stratification approach, where the response of high scoring DS patient can be predicted more accurately that with extant transcriptomic signatures.

**What is already known on this topic:** The predictive power of CTL activity based on bulk tumor transcriptomics, despite being a widely studied important determinant of ICI treatment, is very limited.

**What this study adds:** The efficacy of CTL activity in predicting ICI therapy response is significantly higher among patients with decoupled CTL and ETL activities.

**How this study might affect research, practice or policy:** We identified a set of genes as the decoupling signature, whose upregulation markedly diminishes the correlation between CTL and ETL activities. Our decoupling signature enhances the power of CTL in predicting ICI treatment response, outperforming other extant expression-based signatures.

## Background

Immune checkpoint inhibitors (ICIs) are one of the most important and effective therapeutic strategies that are based on cancer immunology. In 2011, the Food and Drug Administration (FDA) approved ipilimumab (anti-cytotoxic T-lymphocyte-associated protein 4 or anti-CTLA4) inhibitor) as the first ICIs for the treatment of patients with metastatic melanoma^1^. Up to now, there are six more ICIs approved by the FDA, including three anti-programmed cell death protein 1 or anti-PD1 inhibitors: pembrolizumab, nivolumab, cemiplimab-rwlc, and three anti-programmed cell death protein ligand 1 or anti-PDL1 inhibitors: atezolizumab, avelumab, durvalumab. The combination of ipilimumab and nivolumab in the treatment of metastatic melanoma has remarkably improved 5 year overall survival (OS) to 52%^2^, which is an exciting improvement of patient survival compared to the months survival prior to the emerging of ICIs^3,4^. However, about half of melanoma patients still do not benefit from ICI treatment. Thus, there is an unmet need to identify accurate biomarkers to predict which patients will respond to ICIs, and on the longer run, to predict new and even better therapies.

Cytotoxic T lymphocytes (CTL) play a key role in determining patients response to ICIs^5,6^. CTL activity, estimated using signature genes specifically expressed on CTLs^7,8^, has been frequently used to predict ICIs response. However, its prediction power is limited^9,10^. One possible reason is that CTL encounter exhaustion due to the constant stimulation of cancer cells and immunosuppression within the tumor microenvironment (TME)^11,12,13^, mediated via elevating inhibitory receptors (IRs) expression^11^. Such exhausted T cells are characterized by the loss of effector functions, elevated and sustained expression of inhibitory receptors^11^. The emergence of immune checkpoint blocking therapy as a strategy to treat cancer is based on the ability of mAbs to block the interaction between specific IRs on exhausted T lymphocytes (ETL) and their corresponding ligands on cancer and other antigen-presenting cells^14^. Blocking such inhibitory interactions promotes the expansion and recovery of effector function in ETLs, leading to tumor regression in cancer patients. However, the dysfunctional state of terminal exhausted T cell cannot be reprogrammed by ICIs^15,16^. Recent studies reported that majority of tumor neo-antigen specific CD8 T cells were in a state of exhaustion^17,18^.

Analyzing tumor TCGA bulk expression data, we observed a strong positive correlation between CTL and ETL activities (estimated based on conventional gene signatures in the bulk tumor expression) across multiple cancer types. We hypothesized that the limited predictive power of CTL activity might result from the high concordance of CTL and ETL activities in bulk expression, such that their activities mutually cancel out their antagonistic effects on ICI response. Our analysis primarily focuses on bulk data, as the positive correlation between CTL and ETL was specifically observed in bulk data, rather than in single cell data. Consequently, we have set out to develop a computational framework aimed at identifying a set of genes whose expression can identify a subset of patients where the CTL and ETL correlation is diminished as much as possible. We show that, indeed, the power of CTL activity in predicting ICI response in melanoma is enhanced in such patients.

## METHODS

### Transcriptomic data and gene signatures

Gene expression data of TCGA patients were downloaded from GDC: https://portal.gdc.cancer.gov. Transcriptomic data, cell type abundances, and cell-type specific expression profile of ICI treated melanoma cohorts^19–21^ were retrieved from CODEFACS^22^. The following published gene signatures were obtained from original publications: cytotoxic T lymphocytes (CTL) signature^8^, exhausted T lymphocytes (ETL) signature^16^, reactive T cell signature^23^.

### Prediction of ICIs response, patient cohort and clinical end points

Response status of ICIs treated melanoma patients based on the RECIST criteria were retrieved from original publications^19–21^. ‘‘CR/PR’’ patients were classified as responders and ‘‘SD/PD’’ patients were classified as non-responders. Previously published biomarkers, TIDE^7^, IMPRES^,24^, CD274 (PDL1)^25^, stroma EMT^26^, CD8 T cell effector^27^, and TGFB^28^, were collected from the literature and tested for association with response to ICIs therapy. Sample-wise scores were calculated from bulk RNA-seq data using TPM values and following the methodology described in corresponding studies. Genes with unavailable expression data were excluded from calculations of gene signature scores. The predictive utility of these immune signatures was evaluated with AUC values derived from ROC curves of gene signature scores and odd ratio calculated from confusion matrix. Cutoffs for determining responders and non-responders were optimized by maximizing the sum of specificity and sensitivity.

### Identification of decoupling signatures

To identify genes that can mitigate the correlation between CTL and ETL activities. We employed a variable interaction test in a multivariate linear regression to TCGA SKCM cohort.

For each gene *G*, we performed the following regression:

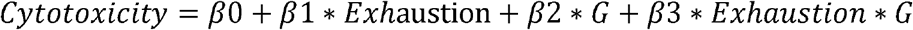

Where *Cytotoxicity* and *Exhaustion* denote the CTL and ETL activity levels which were estimated by calculating the enrichment score for CTL and ETL signatures. *G* represents the expression level of gene *G* in a tumor.

To further understand the variable interaction test, we can rewrite the model as follows:

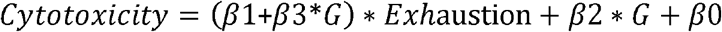

The association between CTL and ETL activities is (*β*1+*β*3*G). The coefficient *β*1 is typically positive because the CTL and ETL activities are positively correlated (**Figure 1A**). The expression value used in this model are positive values, which means a positive coefficient *β*3 will reduce the positive association between CTL and ETL activities, whereas a negative coefficient will enhance the positive association. The decoupling signatures were identified according to the *t* value: *β*3 / StdErr(*β*3) and FDR. A total of 235 genes had significant positive coefficient values and were identified as *decoupling signature* using cutoff T value < 0 and FDR <= 0.01.

**Figure 1.**
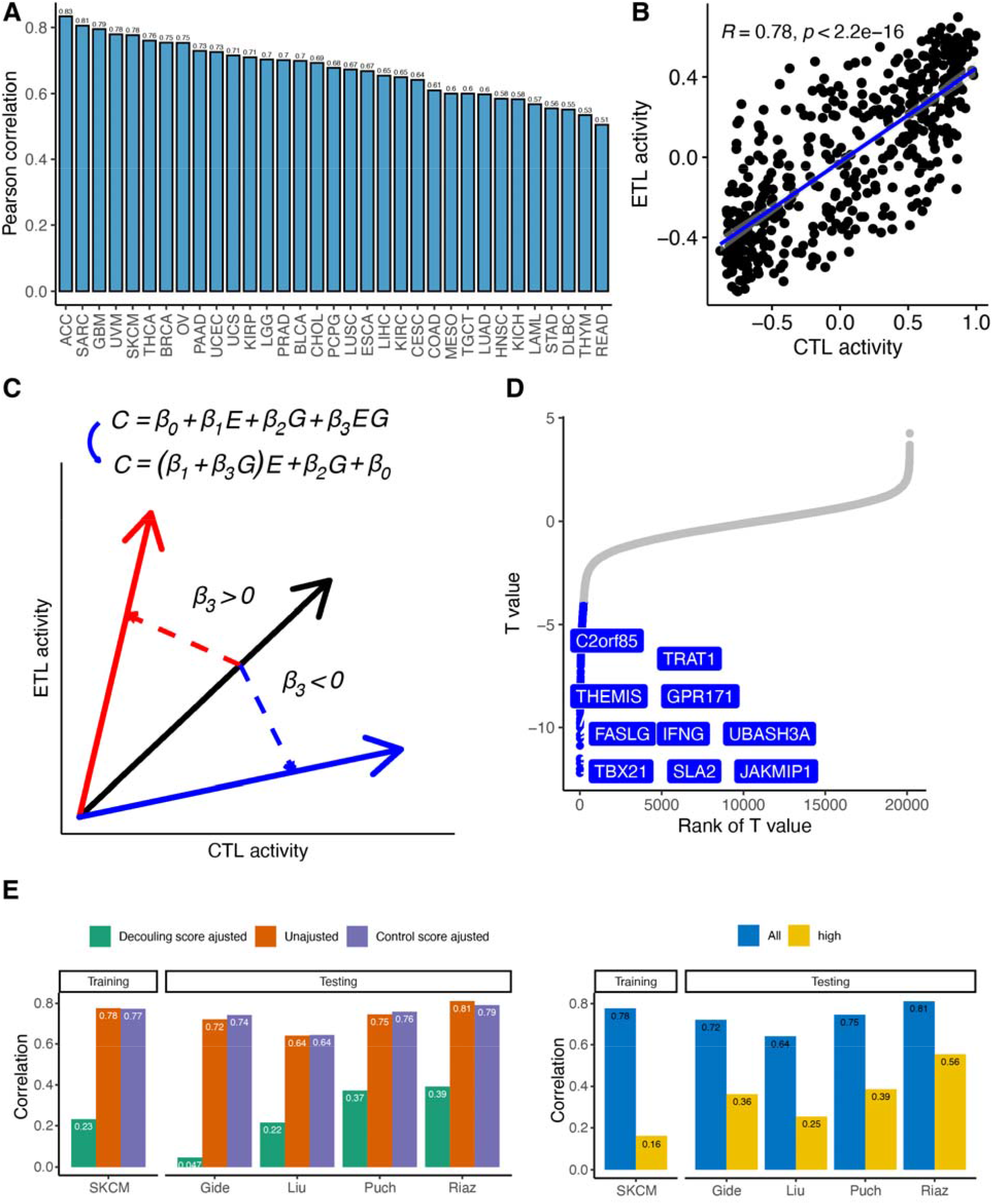
Characterization and decoupling of CTL and ETL activities. **A**. The Pearson correlation between CTL and ETL activities, shown for many different cancer types in the TCGA cohort. **B**. A sample-wise scatter plot of CTL and ETL activities in TCGA Skin Cutaneous Melanoma (SKCM) patients. **C**. The Interaction multivariate linear regression used to identify the decoupling signature set of genes and is applied to all genes, one by one. Variables *E* and *G* represent the relation between T cell exhausted activity and expression of a given candidate gene, respectively. The coefficient of covariate (*E * G)* represents the conjunct interaction effect between *E* and *G* variables activity and CTL activity. The slopes between T cell cytotoxic and exhausted activity are shown by the arrow lines. Since the gene expressions used in this model are positive values, the change of the slope reflects the value of the coefficient *β*3. D. The distribution of significance T values of all genes; those included in the decoupling signatures are colored in blue and a few top ones are specified; a negative T value of the covariate (*E * G)* for all tested genes indicates that the increased expression of this gene can mitigate the correlation between T cell cytotoxic and exhausted activity; if this effect is significant (Methods) it is included in the decoupling signature set. **E**. Decoupling score adjusted correlations between CTL and ETL activities versus the observed when using the random control signatures or the unadjusted (non-decoupled) correlations. **F**. CTL and ETL activities correlations in the high decoupling-score patients’ group (top 20 percent decoupling scores).

### TCGA Tumor Infiltrated Lymphocyte (TIL) patterns

Tumor Infiltrated Lymphocyte (TIL) patterns for TCGA SKCM patients were retrieved from a study^29^. Briefly, pathology sides were used to characterize the TILs patterns for TCGA patients. TILs patterns were visually assigned by a pathologist into one of five categories:

1. Brisk, diffuse: diffused TILs scattered throughout at least 30% of the area of the tumor.
2. Brisk, band-like: band-like boundaries formed by TILs bordering the tumor at its periphery.
3. Non-brisk, multi-focal: loosely scattered TILs present in less than 30% but more than 5% of the area of the tumor.
4. Non-brisk, focal: loosely scattered TILs present less than 5% but greater than 1% of the area of the tumor.
5. None: few TILs were present involving 1% or less of the area of the tumor.

### Statistics and enrichment analysis

Differences between two-continuous variable were assessed using Wilcoxon rank sum test^30^. ANOVA test^31^ was used to determine if there is a statistically significant difference between more than two categorical groups. Association between two-continuous variable were measured by Pearson correlation^32^. To make correlations between groups comparable, we down sampled the majority class(es) in each comparison. R package “ppcor” was used to eliminate the effect of other variables when assessing the correlation between CTL and ETL activities^33^. Gene set enrichment analysis of decoupling signature genes was conducted against the GO BP database using clusterProfiler^34^ with the follow settings: OrgDb = org.Hs.eg.db, ont = “MF”, pAdjustMethod = “BH”, pvalueCutoff = 0.01.

## RESULTS

### Identifying a decoupling signature of genes whose upregulation identifies a subset of melanoma patients with reduced correlation between cytotoxic and exhausted T cell activities

To systemically characterize the association between CTL and ETL activities in bulk RNAseq data, we used established gene signatures of CTL^35^ and ETL^16^ activities. Given these signatures, We analyzed the bulk RNAseq data via ssGSEA algorithm^36^ to estimate CTL and ETL activities in TCGA patients. By calculating the correlation between CTL and ETL activities across all samples in each cancer type, we found that the CTL and ETL activities were highly positively correlated in all cancer types (mean Pearson correlation: 0.67 ± 0.086) (**Figure 1A, Supplementary Fig. 1**). We hypothesized that this correlation may cancel out their opposing associations with ICI response and blunt the predictive power of ETL activity.

As melanoma has rich transcriptomic datasets of patients treated with ICIs, we decided to focus our study on the latter cancer type. In addition, melanoma has a few published transcriptomics biomarker ICI signatures that we can compare with. To decouple the CTL and ETL CD8+ T cell states, we analyzed the TCGA melanoma expression dataset via an interaction linear regression model aimed at identifying genes whose expression status mitigates the positive correlation between CTL and ETL activities (the scatter plot of the two signatures activities across all melanoma samples is shown in **Figure 1B**). In this model, the association between CTL and ETL activities can be represented by the coefficient denoting the slope of CTL and ETL activities (**Figure 1C**). Since the gene expression activities used in this model are positive values, the change in slope depends on the coefficients of the covariates^37^. A positive coefficient value indicates that higher expression of this gene increases the association of CTL and ETL activities, conversely, a negative value indicates that the higher expression of this gene reduces the association. Using this model, we identified a gene set of 235 genes (**Figure 1D**) (**Supplementary file 1**), whose increased expression decouples CTL and ETL activities. We termed this gene set *the decoupling signature*. Following, in each individual tumor sample, its *decoupling score* denotes the enrichment score of the decoupling signature, calculated using single samples Gene Set Enrichment Analysis algorithm (ssGSEA)^36^. Among the decoupling signature genes, some effector and memory T cell-associated genes^38–40^ ranked in the top, such as *IFNG, TBX21, GZMH*. The expression of these genes in the patients indicates that the CD8 T cells in the TME were still functional and had not entered a dysfunctional state.

To verify the capability of the decoupling signature to decouple of CTL and ETL activities, we first calculated the decoupling scores for TCGA melanoma patients (labeled SKCM) and for three independent cohorts ICIs-treated melanoma patients (Gide^19^, Liu^20^, Riaz^21^, Cui^41^). We randomly selected the same number of genes as in the decoupling signature to serve as random control, comparison signatures (Methods). We then employed a partial correlation analysis to compute the correlation between CTL and ETL activities after adjusting for the effect of the decoupling signature and that of a given control signature. Then, we compared both correlations to that of the original correlation (unadjusted) between the cytolytic and exhausted signatures (**Figure 1E**). As expected, the correlation of CTL and ETL activities is indeed markedly decreased after adjusting for the decoupling score compared to that observed after adjusting to the control scores or to the original, unadjusted correlation (**Figure 1E**). We then grouped patients into those with a high-decoupling-score (top 20% patients group). Threshold for the group was defined according to two rules: 1. Include as many patients as possible in the high-decoupling-score group; 2. Minimize the correlation between CTL and ETL activity in the high-decoupling-score group. We tested several thresholds and found that the 20% threshold fits our rule best (**Supplementary Fig. 2A**). In both the training dataset (TCGA melanoma) and testing datasets (ICIs-treated melanoma), the high-decoupling-score groups show a markedly lower correlation between CTL and ETL activities (**Figure 1F**). These results testify to the TCGA-inferred decoupling score ability identify sub-groups of patients where the correlation between CTLs and ETL activities in ICIs treatment melanoma cohorts is diminished.

### CTL activity is highly predictive of ICI response in high decoupling-score melanoma patients

We turned to assess the performance of CTL activity in predicting ICIs response in the high-decoupling-score patients (top 20% patients group). We measured the prediction performance of CTL activity in three independent ICI cohorts using two complementary standard measures, the Area Under the Receiver Operating Curve (AUC) and the Odds ratio of responders to non-responders (OR). The AUC is a standard measure in machine learning for evaluating the overall predictive performance of a classifier across all possible decision thresholds. The OR denotes the odds to respond when the treatment is recommended divided by the odds when the treatment is not recommended. It quantifies the performance at a chosen decision threshold and is hence a more clinically oriented measure (Methods). In 5 of the 6 treatment groups we studied, CTL activity achieved better performance in the prediction of ICIs response in the high-decoupling-score (**Figure 2A, B**). We next compared the CTL activity-based prediction in the high decoupling score patient group to that observed with other contemporary transcriptomics-based ICI response prediction methods and biomarkers, TIDE^7^, IMPRES^,24^, CD274 (PDL1)^25^, stroma EMT^26^, CD8 T cell effector^27^, and TGFB^28^ (Methods). We find that CTL activity exhibits a substantially superior performance compared to other methods and biomarkers. However, it is important to acknowledge that not all of the observed differences reached statistical significance, likely due to the limited sample size (**Figure 2C, D**). These results thus put forward a possible two stage approach for stratifying melanoma patients to ICI treatment from bulk tumor expression: (1) First, identify whether a patient has a high decoupling score, and if so, (2) Second, predict her or his response to ICI treatment by the CTL score computed from her or his tumor expression..

**Figure 2.**
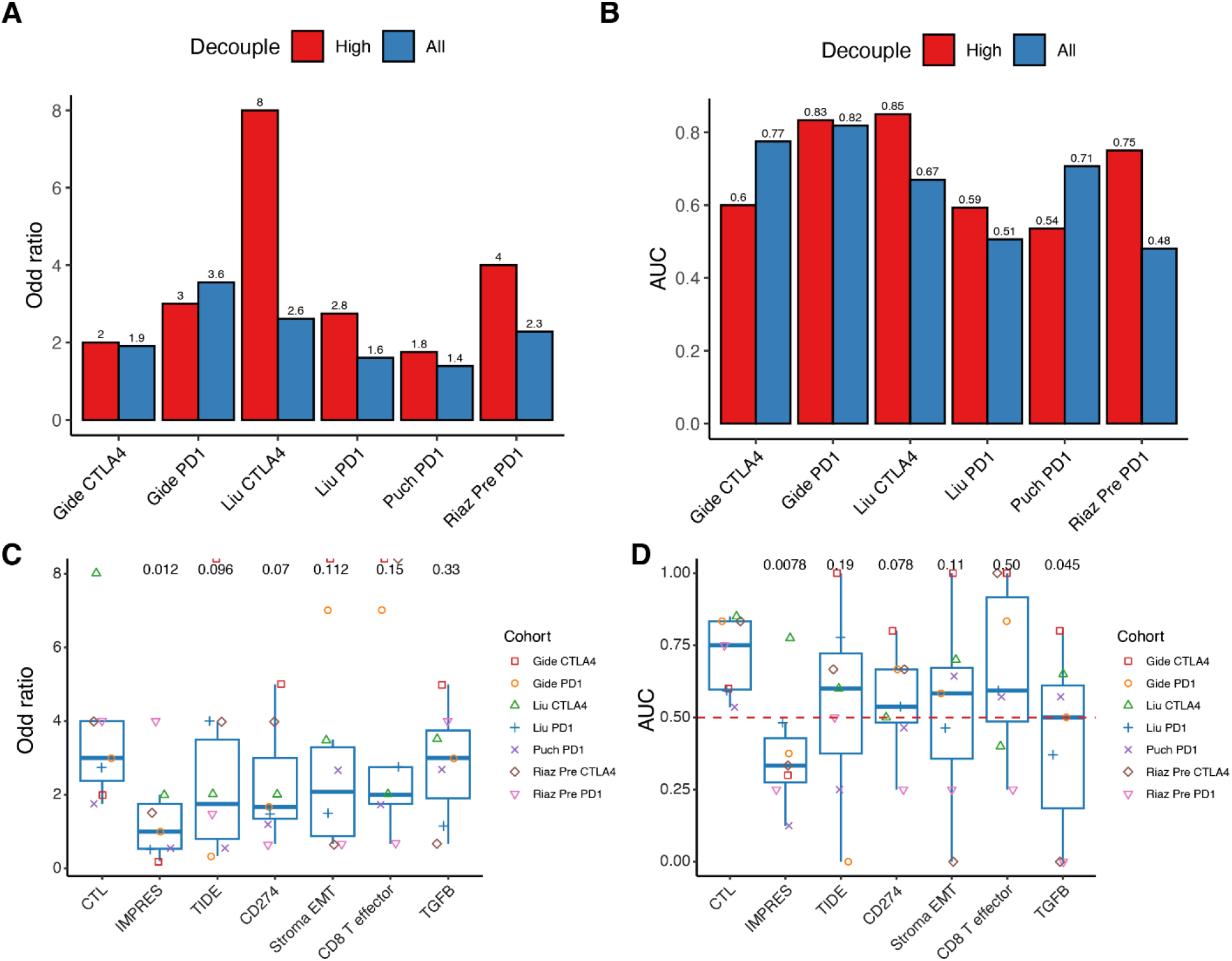
CTL activity is strongly predictive of ICI response in high-decoupling-score patients. Bar plots showing the cytotoxic activity ICI’s response prediction accuracy in different melanoma cohorts. This is shown in terms of both odds ratio (**A**) and Area under the ROC Curve (AUC, **B**) in the high decoupling groups compared to the groups that include all patients. The Riaz Pre CTLA4 treatment group of patients with high decoupling scores was excluded from this analysis as it does not include any non-responders. The odds ratio (C) and AUC (D) quantifying ICIs prediction performance of CTL to the previously established transcriptomics-based signatures in the high-decoupling-score patients. One-sided p-values were displayed at the top of each box and were calculated using a paired Wilcoxon rank test to compare the control (CTL) group with other signatures. In all panels the four ICIs-treated melanoma cohorts studied are sub-divided into seven cohorts according to the treatments the patients have received.

### Uniformly decoupled CTL and ETL activity is predictive of melanoma patients response to ICI therapy

In the previous section, we have shown that cohort-specific percentile-based thresholds can be used to identify a high scoring group of patients where CTL activity is strongly predictive of response. We next aimed to investigate if we can identify two fixed-values thresholds that can achieve these goals across all patients studied, laying a more solid basis for the potential future utilization of this approach in the clinic.

To identify such ‘global’ uniform thresholds, we needed to integrate the decoupling scores of patients from all three ICI-treated melanoma cohorts. Although we processed transcriptome data for these three cohorts using a consistent pipeline starting from raw sequence data (Fastq files), there are remaining strong batch effects between these cohorts that needed to be overcome to this end (**Figure 3A**). The distribution of these decoupling scores varied considerably, making them incomparable between the cohorts as is (**Figure 3B**). We hence applied ComBat^42^ to the patients transcriptome data, which has successfully eliminated batch effects between cohorts (**Figure 3C**). As a result, the distributions of decoupling scores calculated using the batch-corrected transcriptome data is more consistent across these cohorts (**Figure 3D**).

**Figure 3.**
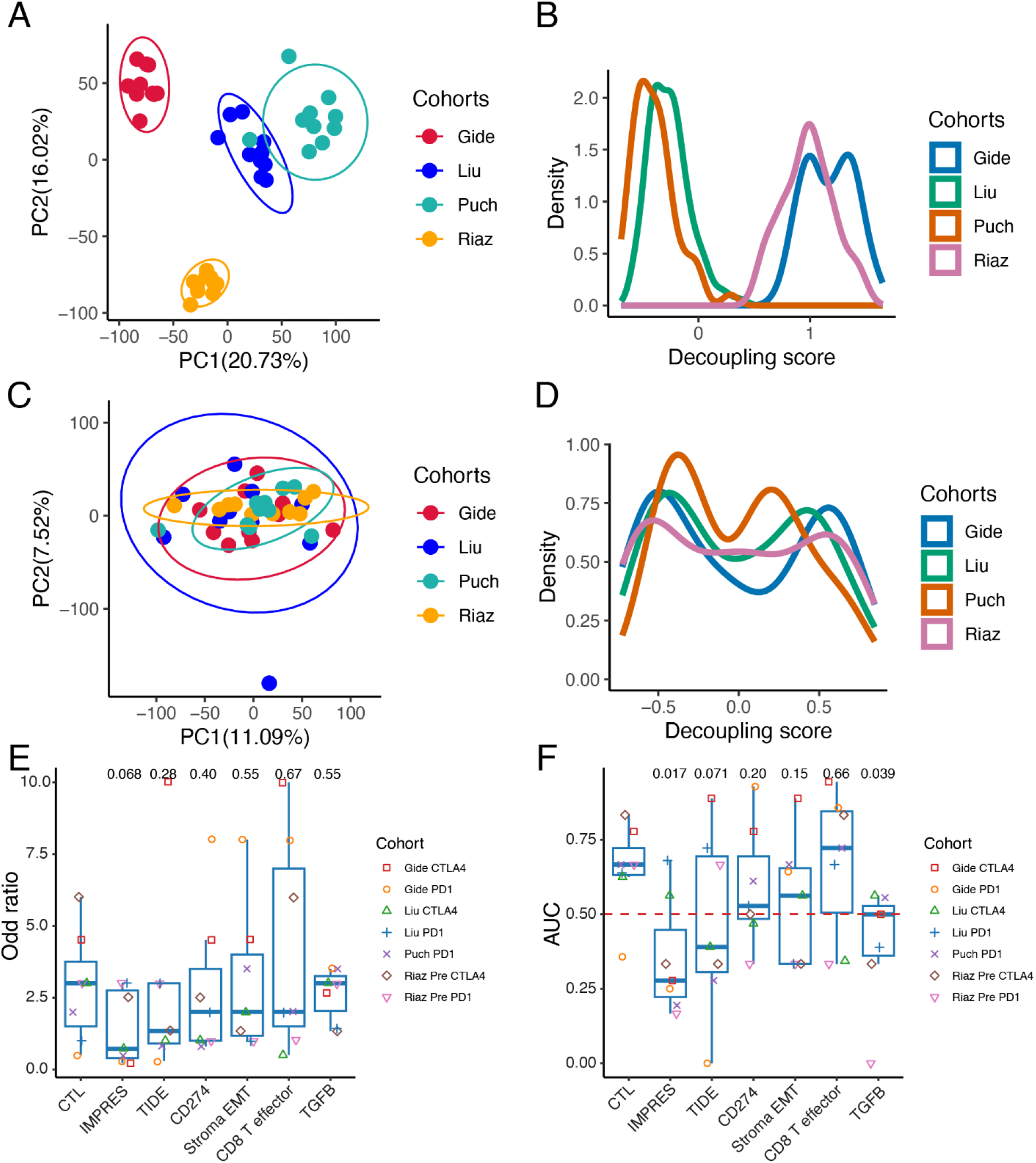
Using uniform high/low thresholds maintains the predictive power of the approach presented. **A**. A PCA plot showing batch effects on transcriptomic data from three ICIs treated melanoma cohorts. **B**. Distributions of decoupling scores computed using transcriptomic data before batch removal. **C**. The PCA plot of transcriptomic data after batch removal. **D**. Distributions of decoupling scores computed using transcriptomic data after batch removal. The odds ratio **(E)** and AUC **(F)** quantifying ICIs prediction performance of CTL compared to that obtained with the previously established transcriptomics-based signatures in the high-decoupling-score patients. One-sided p-values were displayed at the top of each box and were calculated using a paired Wilcoxon rank test to compare the control (CTL) group with other signatures. In all panels the four ICIs-treated melanoma cohorts studied are sub-divided into seven cohorts according to the treatments the patients have received.

Given the batch-corrected decoupling scores in all three cohorts, we took the mean of the 20% percentile based high and low decoupling groups thresholds in each cohort as two uniforms high and low thresholds, which are then used to determine the patients in the respective high and low groups across all patients. (**Supplementary Fig. 3A**). These global thresholds identify an about similar number of patients in the high and low decoupling groups across cohorts (**Supplementary Fig. 3B**). We then compared the response prediction performance of ICIs for CTL activity to other transcriptomics-based state-of-art ICI response prediction methods, CTL activity still generally retains its power for predicting ICI response quite better than the other state-of-the-art transcriptomics-based methods (**Figure 3E, F**). Taken together, these results testify to the feasibility of the two-step approach when using these global thresholds.

### The Decoupling signature activity is strongly associated with T cell activation and immune cell infiltration in melanoma

To further characterize the biological pathways contributing to the decoupling signature, we identified the pathways enriched in these genes (Methods). Those point to T cell activation/differentiation and lymphocyte activation and differentiation (**Figure 4A**). Furthermore, the decoupling scores are positively correlated with computationally estimated T CD8, T CD4, macrophage, B cell abundances in the TCGA cohort and in the three melanoma ICI treatment cohorts we studied (Methods, **Figure 4B**). Notably, they are anti-correlated with abundance of tumor cells (a correlate of tumor purity).

**Figure 4.**
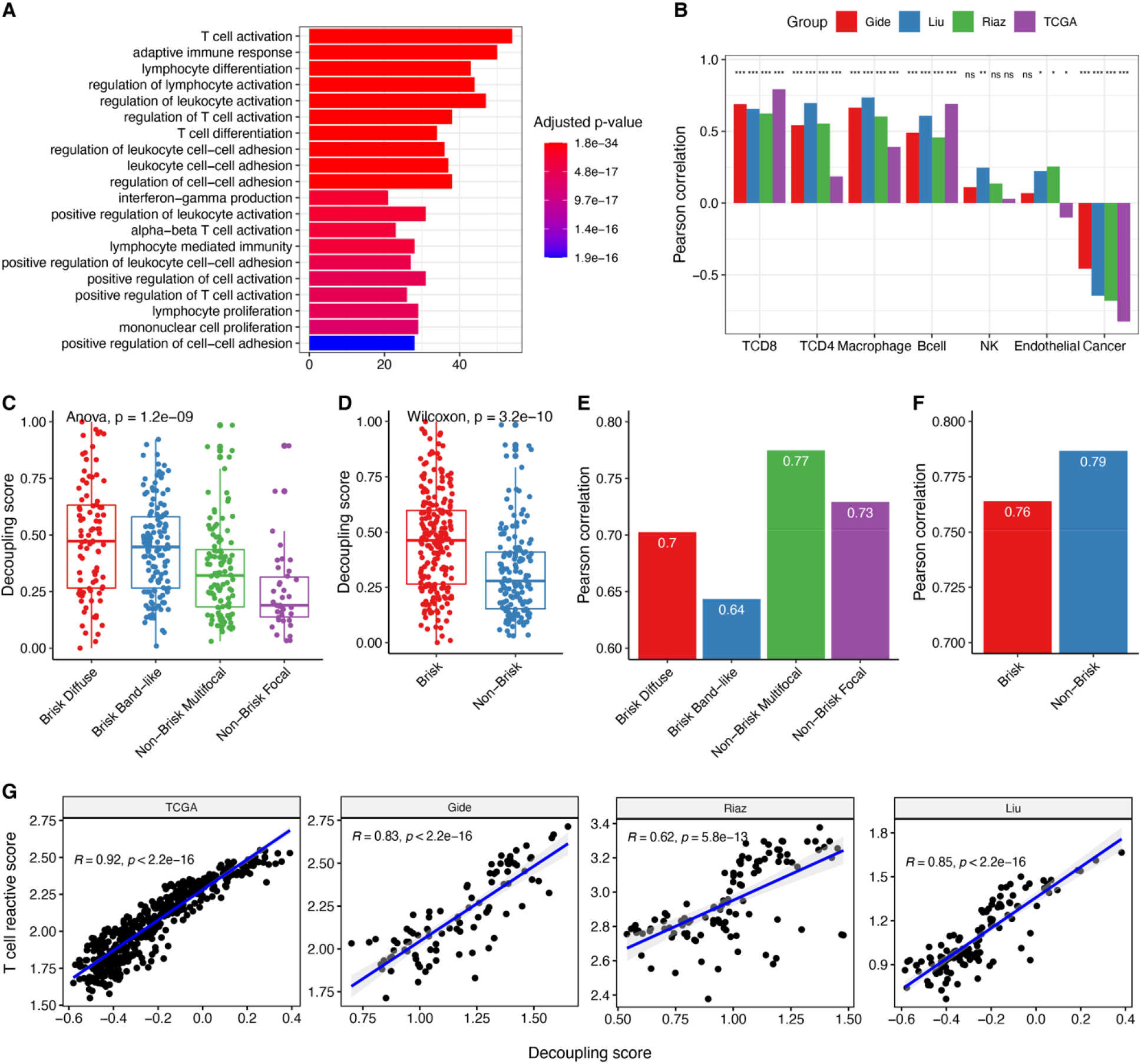
Decoupling signature activity is associated with T cell activation, immune cell infiltration and antigen-specific reactive T cell activity. **A**. Pathway enrichment analysis of decoupling signature genes; the x-axis represents the number of decoupling signature genes in each enriched pathway, the colors represent the corresponding enrichment hypergeometric statistical significance, as indicated in the color schema on the right side (Methods). **B**. The correlation between the decoupling score and cell abundances across TCGA SKCM and three ICIs treatment cohorts; the stars at the top denote significance values (Methods). **C**. The distribution of decoupling scores among patients with different pathology-determined lymphocyte infiltration status. **D**. The distribution of decoupling scores in Brisk versus Non-Brisk tumors. **E**. The correlation between CTL and ETL activities in groups of tumors having a different lymphocyte infiltration status. **F**. The correlation between CTL and ETL activities within Brisk and Non-Brisk tumors. **G**. Scatter plots showing the correlation between the decoupling scores and the tumor antigen-specific reactive T cell activity scores in four different melanoma patients cohorts.

We next set out to study to what extent is the decoupling signature genes activity associated with estimates of T cell abundance and infiltration done by pathologists reviewing tumor H&E slides. To test that, we retrieved lymphocyte status annotations of TCGA melanoma patients from a study that assessed lymphocyte status based on pathological slides data^29^. In this study, patients were assigned to one of five patterns based on tumor-infiltrating lymphocyte status: Brisk diffuse, Brisk band-like, Non-brisk multi-focal, Non-brisk focal, and None^29^ (Methods). “Brisk” represents a moderate to strong immune response with a relatively dense and high proportion of TILs in TME, while “Non-brisk” is indicative of a weak immune response with a scattered and low proportion of TILs in TME. Due to its limited sample size, we filtered the “None” group from further analysis. By comparing the distribution of decoupling scores between samples in the different patterns, we find that the decoupling scores are positively associated with tumor-infiltrating lymphocytes abundance and infiltration in melanoma (**Figure 4C**). We then classified patients dichotomously into Brisk (Brisk diffuse, Brisk Band-like) and Non-brisk tumors (Non-brisk multi-focal, Non-brisk focus, None). Again, we find that Brisk tumors have significantly higher decoupling scores than Non-brisk tumors (**Figure 4D**). Notably, the correlations of CTL and ETL activities in tumors with high infiltration are indeed lower than these correlations in the low infiltration tumors (**Figure 4E, F**). Taking together, these strong associations indicate that in tumors highly infiltrated by T cells, the CTL and ETL activities are relatively lowly correlated, suggesting that the activated CD8 cells are indeed pre-exhausted.

Previous studies have shown that tumor-infiltrating CD8+ T cells consist of virus-specific bystander T cells and tumor antigen-specific reactive T cells^43,44^. A recent study characterized antigen-specific reactive T cells by combining single-cell sequencing and TCR sequencing technologies^23^. To study the association between decoupling scores and tumor antigen-specific reactive T cell activity, we estimated the reactive T cell activity in each patient’s tumor in the melanoma cohorts we study, based on the reactive T cell activity signature of^23^. As the tumor bulk expression is a mixture of all cell types in the tumor microenvironment, we used the deconvoluted gene expression profiles of CD8 T cells from^22^ to estimate the reactive T cell activity in melanoma patients in the different cohorts we have studied. Notably, we find that the decoupling scores are highly positively correlated with the estimated reactive antigen-specific T cell activities (**Figure 4G**). These results suggest that the decoupling score is not only associated with the overall abundance of TILs, but also of with the abundance of tumor antigen-specific reactive T cells. It further supports that the notion that in the subset of tumors were the CTL and ETL activities are not correlated, the tumor environment is indeed enriched with cytolytic non-exhausted and reactive T cells.

## DISCUSSION

Motivated by the strong associations observed between CTL and ETL activities estimated from bulk RNAseq data across cancer types, our objective was to devise a computational framework to disentangle their respective contributions. We show that the resulting decoupling signature, inferred from TCGA ICIs treatment-naïve bulk RNAseq melanoma data, can successfully decouple CTL and ETL activities in melanoma. Notably, this decoupling signature can be used to enhance the ability to use CTL activity to predict the response of melanoma patients to immune checkpoint inhibitors in a subset of high-scoring patients, outperforming the predictions obtained by using other contemporary transcriptomics-based predictors. Decoupling signature genes are enriched in immune response-related pathways such as T cell activation and differentiation. Furthermore, the decoupling score is positively correlated with lymphocyte infiltration and negatively correlated with tumor purity estimates. These findings suggest that the efficacy of CTL in predicting ICB response depends on the level of T-cell infiltration, as indicated by the level of the decoupling signature.

The results presented here present a conceptually novel approach for stratifying patients to ICI therapy based on the sequential use of two transcriptomic based biomarkers. It presents an obvious limitation, as the tradeoff inherent in such an approach is that one achieves higher predictive accuracy on a limited subset of patients, and yet, as we show, it bears considerable benefits. It would be of interest to study the potential value of the approach presented here to predict ICI response in other cancer types, since as we have seen in many of them the CTL and ETL activities are strongly correlated. Beyond that, we hope that the two-step stratification approach presented here may motivate additional approaches for systematically identifying confounders of existing predictors and further decoupling such confounding effects. This will gradually make ICIs response more predictable and potentially benefit more cancer patients.

We have shown that given a cohort of melanoma patients, the existing CTL signature is quite highly predictive of the response of these patients to therapy in the subset of 20% of patients with the highest decoupling scores. This result has an underlying biological rationale, since as we have shown in these patients CTL activity is decoupled from the ETL activity, eliminating the potentially masking effects of activated T cells that are essentially also exhausted. Going beyond that, we also have shown that one can use the double-pronged approach to predict patients response in patients whose decoupling scores are higher than a given predetermined threshold value. The latter result lays the foundation for considering the use of this approach to predict the response of individual patients arriving in a newly formed clinical trial cohort. We hope that the results presented here may help facilitate such future studies.

## List of Abbreviations

CTL: Cytotoxic T lymphocytes
ETL: Terminal exhausted T lymphocytes
DS: Decoupling signature
ICIs: immune checkpoint inhibitors
TME: tumor microenvironment
OS: Overall survival
IRs: Inhibitory receptors

## Acknowledgments

This research was supported in part by the NIH Intramural Research Program, National Cancer Institute. This work utilized the computational resources of the NIH HPC Biowulf cluster. (http://hpc.nih.gov).

## Supplemental Material

**Supplementary Figure 1.**
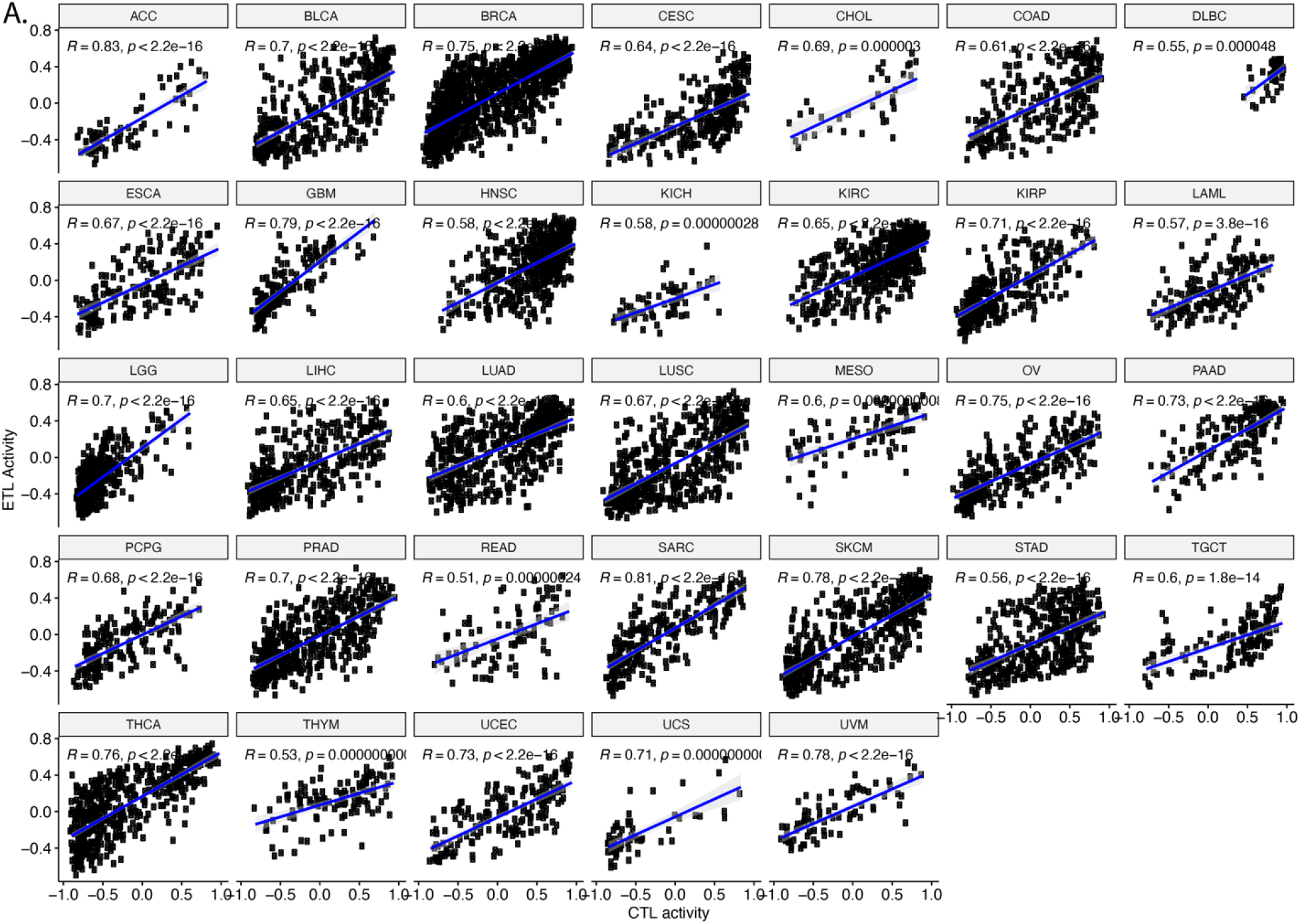
Scatter plots showing the Correlation of CTL and ETL activities in different cancer types in the TCGA cohort. **A**. Each dot represents a patient. The X axis and Y axis represent CTL and ETL activities estimated using the corresponding gene signatures as described in the main text.

**Supplementary Figure 2.**
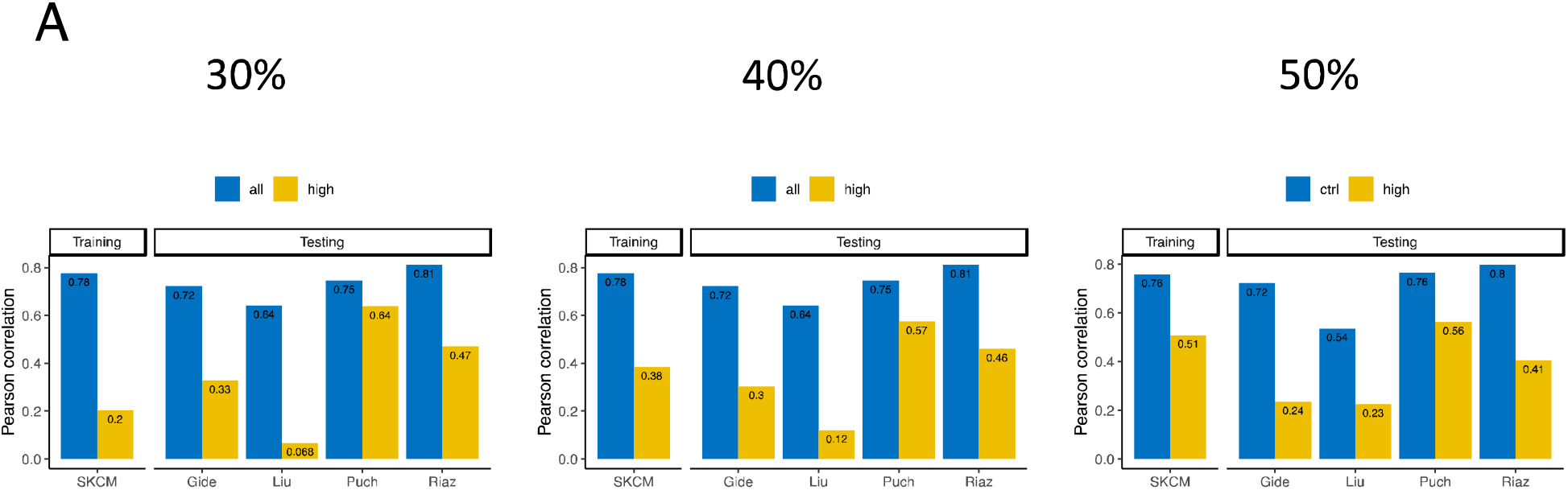
The decoupling scores obtained at different thresholds of high vs low percentile groups. **A**. CTL and ETL activities correlations in the high-decoupling-score patients’ group and the low-decoupling-score group. As evident, the 20% threshold achieves the best middle ground between achieving high decoupling scores while still covering a sizeable fraction of the patients.

**Supplementary Figure 3.**
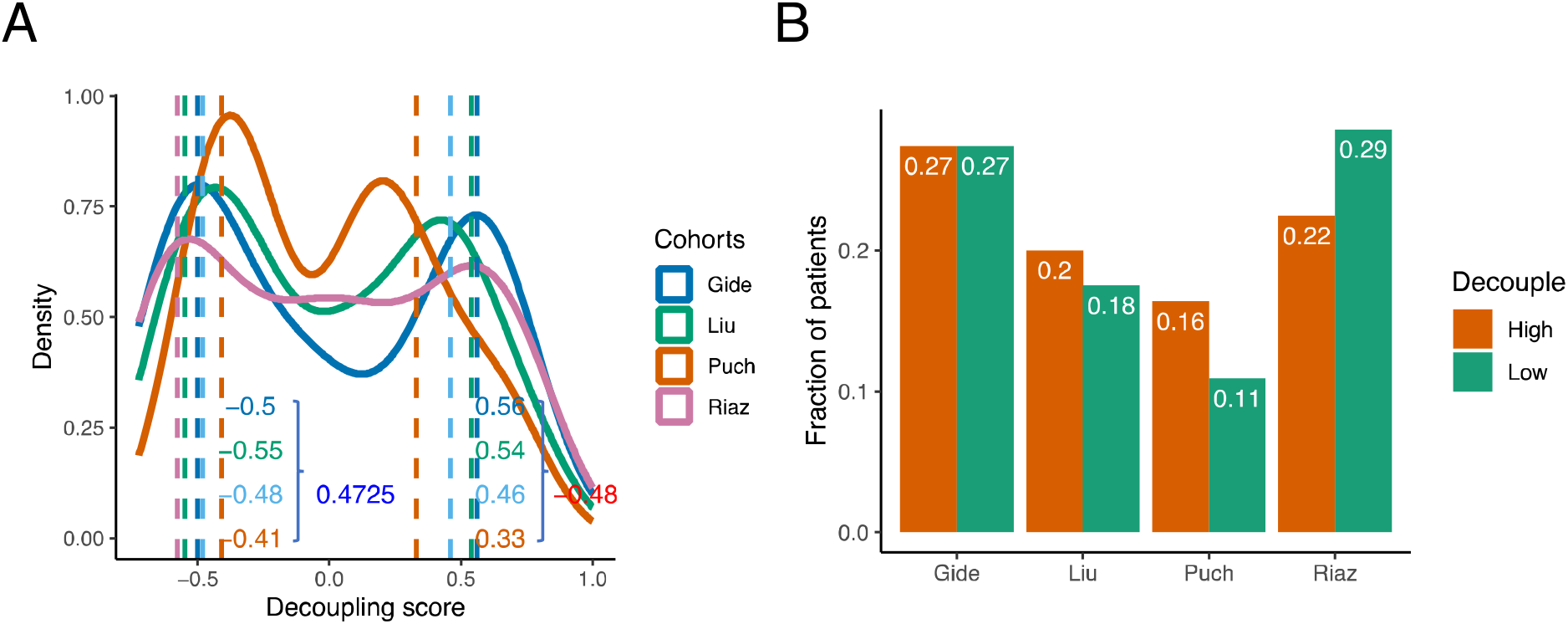
Identification of the uniform thresholds determining the high and low decoupling scores groups of patients in the ICI cohorts. **A**. The solid lines represent the distributions of decoupling scores in each of the three ICIs treated melanoma cohorts. Dashed lines represent the cutoffs of 20% highest decoupling score patients and 20% lowest decoupling score patients in each cohort. The values of the global uniform high and low thresholds on the decoupling scores were determined as the mean of the 20% respective thresholds determined individually in each cohort. **B**. The fractions of patients in the high and low decoupling score groups in each cohort, as determined by using the respective high and low uniform thresholds determined in A.

## Notes

### Competing Interest Statement

E.R. is a co-founder of MedAware, Metabomed and Pangea Biomed (divested), and an unpaid member of Pangea Biomed's scientific advisory board. The other authors have no competing interests.

### Summary of Updates

Update figures and authors

